# Competition between binding sites determines gene expression at low transcription factor concentrations

**DOI:** 10.1101/033753

**Authors:** David van Dijk, Eilon Sharon, Maya Lotan-Pompan, Adina Weinberger, Eran Segal, Lucas B. Carey

## Abstract

The response of gene expression to intra-and extra-cellular cues is largely mediated through changes in the activity of transcription factors (TFs), whose sequence specificities are largely known. However, the rules by which promoters decode the amount of active TF into gene expression are not well understood. Here, we measure the activity of 6500 designed promoters at six different levels of TF activity in budding yeast. We observe that maximum promoter activity is determined by TF activity and not by the number of sites. Surprisingly, the addition of an activator-binding site often reduces expression. A thermodynamic model that incorporates competition between neighboring binding sites for a local pool of TF molecules explains this behavior and accurately predicts both absolute expression and the amount by which addition of a site increases or reduces expression. Taken together, our findings support a model in which neighboring binding sites interact competitively when TF is limiting but otherwise act additively

**Significance Statement:** In response to intracellular and extracellular signals organisms alter the concentration and activity of transcription factors (TFs), proteins that regulate gene expression. However, the molecular mechanisms that determine the response of a target promoter to changes in the number of active TF molecules are not well understood. By combining mathematical modeling with measurements of TF dose-response curves for thousands of designed promoters, we show that competition for active TF molecules is a major factor in determining gene expression. At low TF concentrations additional activator-binding sites within a promoter can actually reduce expression. Thermodynamic modeling suggests that steric hindrance between neighboring binding sites cannot explain this behavior, but that competition for limiting TF molecules can.

## Introduction

Cells respond to internal and external changes by controlling their gene expression programs. A major mechanism by which this is achieved is by modulating the activity of transcription factors (TFs) that bind to specific sites in gene promoters where they activate or repress transcription(Struhl, 1995). For example, in the budding yeast
 *S. cerevisiae*, almost half of the genome changes in expression level in response to amino acid starvation. A single transcription factor, Gcn4, is responsible for the activation of over 500 of these genes(Natarajan et al., 2001). While transcriptome and Chromain-IP (ChIP) studies are useful for understanding the wiring of these large regulatory networks, they are not informative about how the quantitative relationship between TF and target gene expression are encoded in the DNA. It is still not well understood how promoter architecture determines how each target of a TF will respond to changes in the amount of active TF ([TF]). Furthermore, many targets of the same transcription factor are expressed at different levels in the absence of that TF, and the fold-induction of the target is largely independent of its expression at low or high TF (Carey et al., 2013; Rajkumar et al., 2013). However, the molecular mechanisms that enable this decoupling are largely unknown.

In order to understand how promoters encode the function that maps the amount of active TF to a gene transcription level output, we measure the dose response curves for 6500 synthetically designed promoters. We have used a synthetic approach(Sharon et al., 2012) in which pairs of promoters differ by a single regulatory element, in contrast to native promoters that have many differences between them, preventing systematic investigation of the effect of individual DNA sequence elements on expression response.

We observe a wide range of dose response curves that differ in both TF independent expression, and TF dependent (dynamic range) expression. The first, TF independence, can be attributed to differences in the predicted nucleosome occupancy of the promoters. In contrast, TF dependent (dynamic range) differences are mostly determined by the number and affinity of binding sites. Surprisingly, we find that expression level saturates with number of binding sites, at a level determined by the amount of active TF ([TF]) and not by the number of binding sites. Moreover, in many cases adding an activator can actually reduce expression, especially at low [TF]. In order to quantitatively understand the results, we test several hypotheses using a thermodynamic model that incorporates cooperative and competitive interactions between TF binding sites. Our results suggest that expression synergism (i.e. the additive or reductive effect of adding an additional homotypic binding site) is due to ‘sharing’ a local pool of TF between nearby sites.

## Results

### Promoter DNA sequence can encode a wide range of transcriptional responses to changes in the amount of active TF

To systematically measure how transcriptional responses are encoded in promoter DNA sequence we measured the activity of 6500 designed promoters using a fluorescence reporter (Sharon et al., 2012) in six growth media that each differ in their concentration of amino acids ([AA]) (see **Methods** for details). The majority of these promoters contain binding sites for Gcn4, Leu3, Met31 or Bas1; TFs involved in amino acid biosynthesis. At high [AA] the TFs Gcn4, Bas1, Leu3 and Met31 are mostly inactive(Struhl, 1992); at low [AA] the concentration of the active form of these TFs increases (their expression and/or ability to activate transcription increases), and their targets increase in expression(Gasch et al., 2000). For these four TFs, the number of active TF molecules ([TF]) increases gradually in response to decreasing [AA] (Ljungdahl and Daignan-Fornier, 2012) (**Supplementary Fig. 1**). The combinatorial fashion in which TF binding site type, number, affinity, position and accessibility vary in the designed promoter set enables us to systematically investigate the mapping between promoter DNA sequence, [TF] and the induced expression (**Fig. 1A,** see Methods for details).

**Figure 1.**
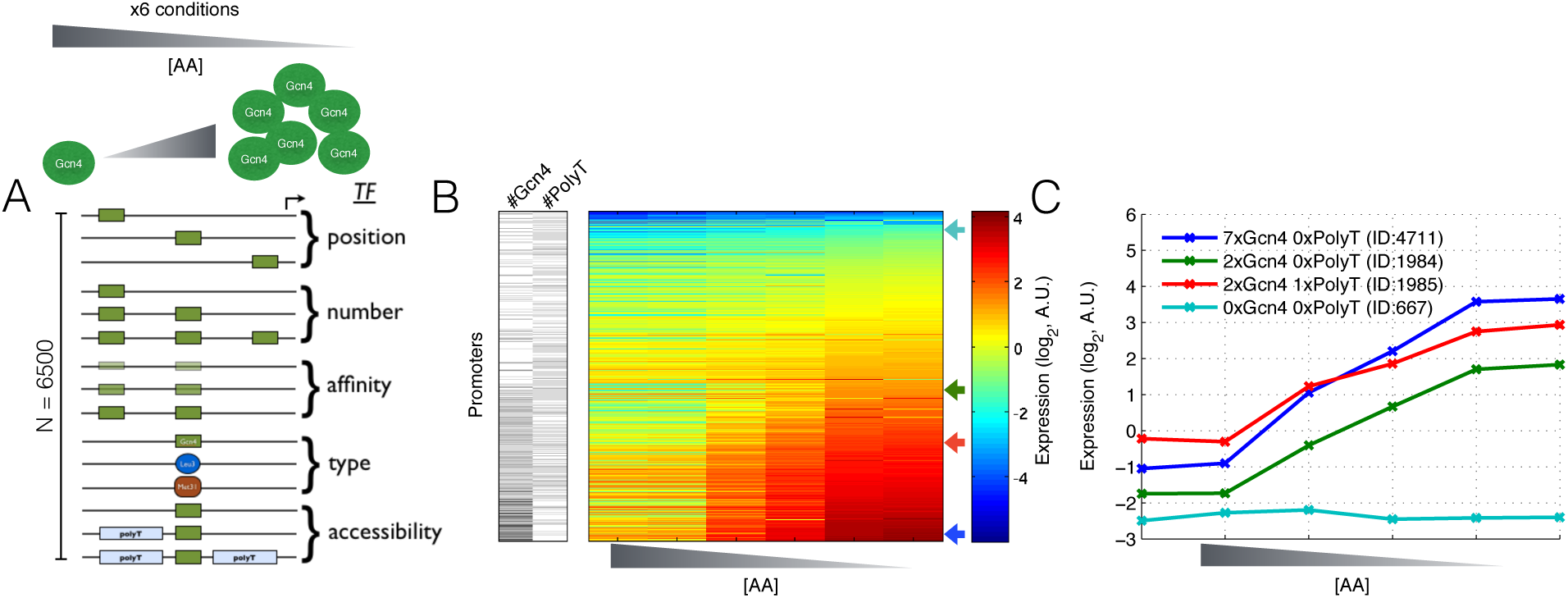
Measurements of TF concentration dependent expression for thousands of designed promoters. **(A)** Schematic depiction of the experimental design. A pooled library of 6500 designed promoters was transformed into yeast and expression levels of all strains in the pooled library were measured in minimal media at each of six different amino acid concentrations (see Methods). **(B)** Promoter expression measurements sorted by dynamic range. For each promoter in the library we obtain an expression measurement at each of the six AA concentrations. For promoters that lack Gcn4, Leu3, Bas1 or Met31 sites, expression does not change with decreasing AA concentration (top of **B**). For promoters with multiple Gcn4 binding sites, expression increases with decreasing [AA]. **(C)** Shown are four representative induction curves showing the effect of changing the number of Gcn4 binding sites (cyan, green, blue), or adding a polyT nucleosome disfavoring sequence (green, red). IDs show library construct identifiers.

The measurements were carried out using our previously described method that involves FACS sorting and deep sequencing of a barcoded pooled promoter library(Sharon et al., 2012; 2014). Briefly, uniquely barcoded promoters that drive a YFP reporter are FACS sorted into 12 bins of expression that subsequently receive an expression-bin barcode. Deep sequencing thus results in reads that contain both a sequence and an expression barcode. A computational analysis of these reads gives, for each promoter and growth condition, an expression distribution, from which the mean is extracted, resulting in 6500 highly reproducible (**Supplementary Fig. 2**) dose-response curves (**Fig. 1B**). Our promoters encode a wide range of responses with a general trend in which more TF binding sites give a greater dynamic range between low and high [AA] (**Fig. 1B**). We observe that some promoter sequence changes (e.g.: addition of a polyT, **Fig. 1C**) affect expression independent of [AA], whereas others (e.g.: addition of Gcn4 binding sites, **Fig. 1C**) affect expression in a manner that depends on [AA]. We refer to the former as (active) [TF] independent expression change, and the latter as [TF] dependent expression change.

### Decoupled control of TF dependent and TF independent expression

In order to distinguish between promoter sequence features that affect expression in a [TF] dependent manner and those that affect expression in a [TF] independent manner, we compare expression at high and low [AA] (see **Methods**) for promoters grouped by DNA sequence features. We find that the number of Gcn4 binding sites affects expression in a TF dependent manner: adding binding sites results, on average, in little increase in expression at high [AA] but a large increase at low [AA] (**Fig. 2A,D**), and thus an increase in the promoter’s dynamic range (**Fig. 2E**). The same results are observed for increasing the affinity of the Gcn4 binding site: increasing the affinity results in slightly higher expression at high [AA], much higher expression at low [AA], and an overall increase in the dynamic range of the promoter (**Fig. 2B,F,G,H**). Thus, both the affinity and number of Gcn4 sites affect a promoter’s expression in a manner that depends on the [TF]. In contrast, adding an additional polyT nucleosome disfavoring sequence results in the same fold change in expression at low and high [AA] and no change in the dynamic range of the promoter (**Fig. 2C,I,J,K**). Adding a binding site for a repressor, changing the position of the binding site, or changing the promoter sequence context to a context with a different predicted nucleosome occupancy also results in no change in the dynamic range (**Supplementary Fig. 3**). Thus, altering the nucleosome occupancy results in a [TF] independent change in expression.

**Figure 2.**
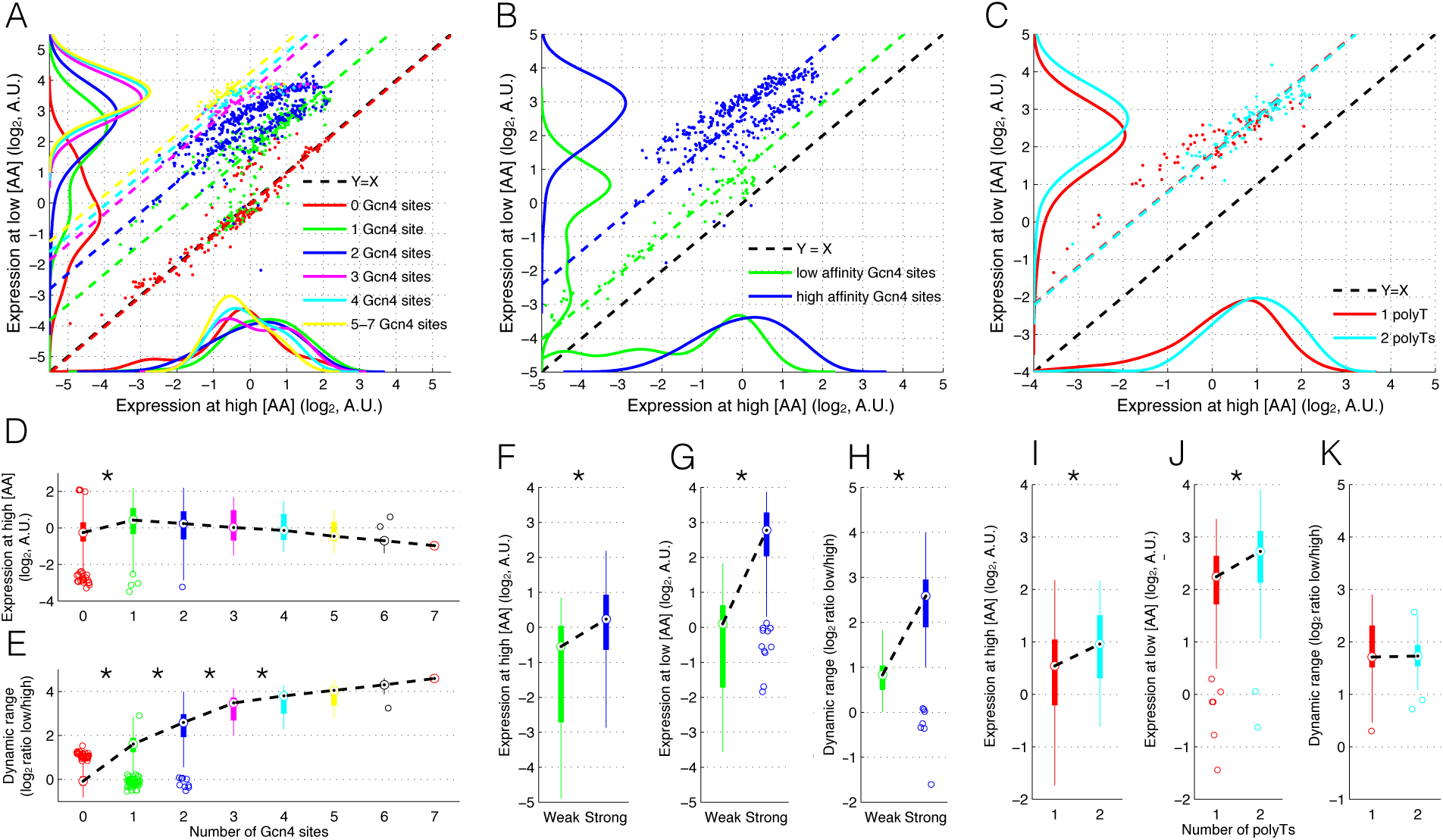
The effect of Gcn4 binding site number and polyT nucleosome disfavoring sequences on [TF] dependent and independent expression. **(A-C)** show expression at high [AA] (X-axis) versus expression at low [AA] (Y-axis) for various promoter sequence features. Dashed lines are the diagonal (slope=1) line that best fit each category of promoters. The black dashed diagonal line (Y=X) represents the regime where expression is constant across conditions. The vertical distance from the Y=X line measures how much any one promoter changes in expression across conditions. Density plots (using ks denstiy estimation) at the X-axis and Y-axis show the distributions of expression values for each promoter at high and low [AA] respectively. **(D-K)** show expression and expression fold change (Y-axis) in box plots as a function of promoter sequence features (X-axis). The dashed black lines in connects the medians of each box. Asterisks denote statistically significant (t-test, p<0.01) changes between subsequent groups. **(A)** Shown are promoters grouped by the number of Gcn4 binding sites. **(B)** Shown are promoters with either low or high affinity Gcn4 sites. **(C)** Shown are promoters with either one or two polyT nucleosome disfavouring sequences. **(D,E)** Box plots of the data in **(A)**. Promoters are grouped by the number of binding sites. **(D)** Shown is expression at high [AA] (Y-axis). **(E)** Shown is expression fold change (dynamic range, log2(low [AA] / [high AA])). **(F-H)** Box plots of the data in **(B)**. Shown are expression at high [AA] **(F)**, low [AA] **(G)** and expression fold change (dynamic range) **(H)** for promoters with low or high affinity binding sites. All differences are statistically significant (t-test p<1e-3, p<1e-5, p<1e-4 for **F**,**G**,**H** respectively). **(I,J,K)** Box plots of the data in **(C)**. Shown are expression at high [AA] **(I)**, low [AA] **(J)** and expression fold change (dynamic range) **(K)** for promoters with either one or two polyT sequences. Expression at high and low [AA] show significant change as a function of polyT number (t-test p<1e-4, p<1e-4 respectively), however dynamic range does not change significantly (t-test p=0.77).

**Figure 3.**
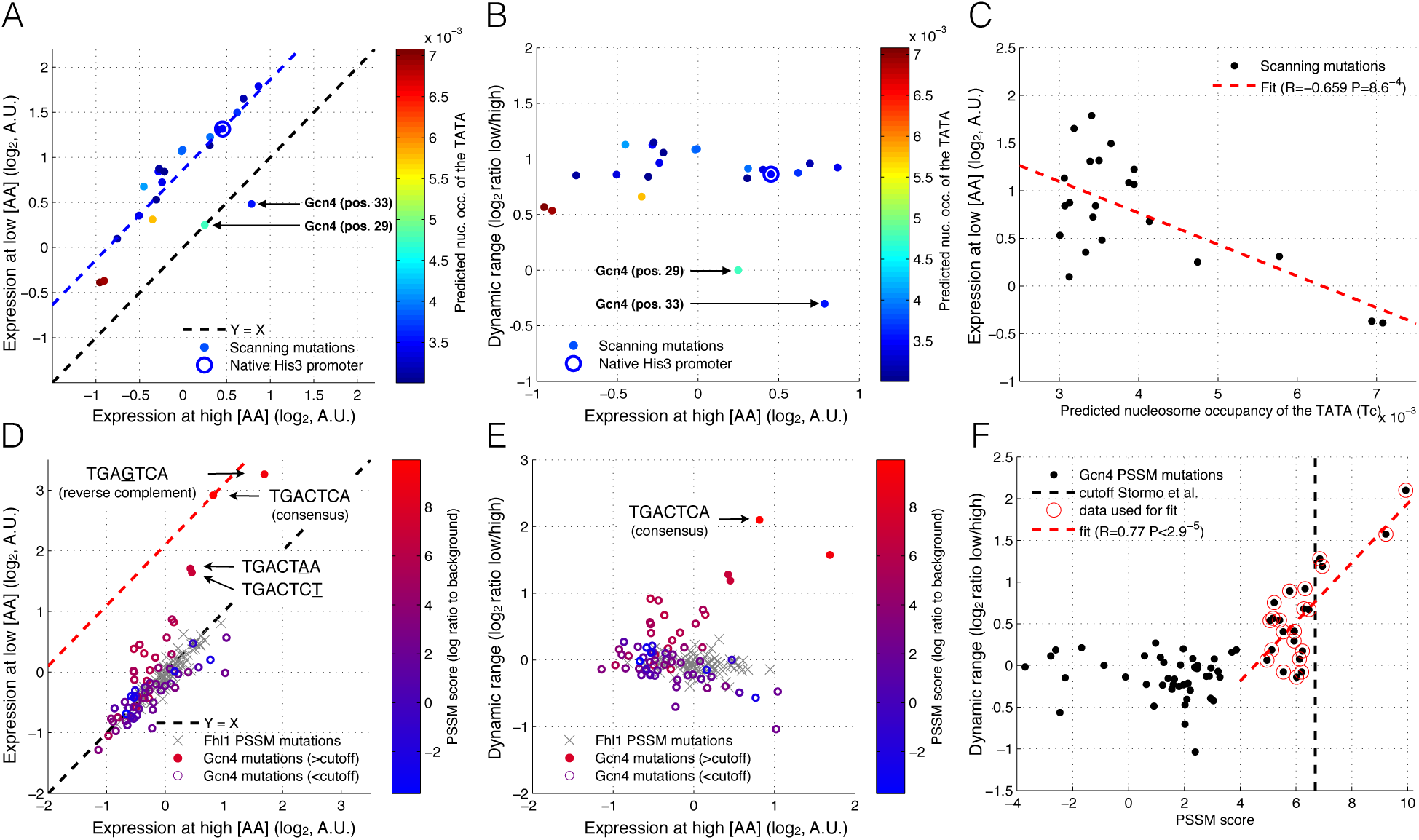
The effect of promoter and binding site mutations on [TF] dependent and independent expression. (A-C) Expression data for a set of 20 sequences that only differ by a single point mutation made at different locations along the promoter. For each sequence the nucleosome occupancy over the TATA box was predicted using a thermodynamic model(Kaplan et al., 2009). **(A)** Shown is the effect of promoter point mutations on expression at low and high AA concentration; the dashed blue line is the best fit of a line with slope=1 to the data. The two scanning mutations that affect the Gcn4 binding site are identified with arrows. These mutations result in removal of the effect of [TF] on expression change. **(B)** The same data as in (A) with dynamic range (log2(low [AA] / [high AA])) graphed against expression at high [AA]. **(C)** Mutations that increase the predicted nucleosome occupancy over the TATA box decrease expression in a linear manner (fit red line). **(D)** A set of promoters, each with a single Gcn4 binding site, with single base mutations that affect the predicted affinity. Arrows mark the consensus sequence and the native His3 promoter sequence. Point mutations in a set of promoters with a single Fhl1 binding site are shown as a control. The dashed red line (slope=1) marks the sequence that gives the highest dynamic range in expression. **(E)** The same data as in (D) with dynamic range graphed against expression at high [AA]. **(F)** Mutations predicted to have a high Gcn4 binding site affinity have a high dynamic range (dashed red line), but the PSSM has less predictive power at low scores.

Taken together, these results show that sequence mediated expression changes affect the dynamic range of expression when they change binding site affinity or number, but not binding site accessibility, and that both the TF dependent and independent behavior of promoters can be tuned separately.

### Mutations inside binding sites affect expression in a TF dependent manner

While it is intriguing that addition or removal of entire promoter sequence elements can alter expression either in a TF dependent or independent manner, we wondered if the same independent control can be achieved by single point mutations that are more readily available in an evolutionary context. To determine this, we examined a set of 213bp scanning mutations made every 3bp across the native HIS3 promoter. We find that 19 of these affect expression in a TF independent manner (t-test p=0.83, **Fig. 3A,B**) and that mutations that increase the predicted nucleosome occupancy over the TATA box have lower expression (Pearson R=-0.66 p=9*10^−4^, **Fig. 3C**). Two of the mutations, which fall within the native Gcn4 binding site, appeared to effectively remove response to [AA] change.

In addition, we find that systematically mutating the Gcn4 binding site results in a change in dynamic range that is correlated with PSSM score (**Fig. 3D–F, Supplementary Fig. 4**). We observe a relatively small increase in expression at high [AA] (Pearson R=0.20, p=0.09), and a much larger increase in expression at low [AA] (Pearson R=0.52 p<3*10^−6^), resulting in a net increase in dynamic range with increasing PSSM score (Pearson R=0.63 p<1.3*10^−8^ or R=0.77 p<3*10^−5^ when only including values above a previously determined cutoff (Spivak and Stormo, 2012), **Fig. 3E,F, Supplementary Fig. 4C**). These results are consistent with models in which low-affinity binding sites are always functional, but have a more pronounced effect at high [TF](Carey et al., 2013).

**Figure 4.**
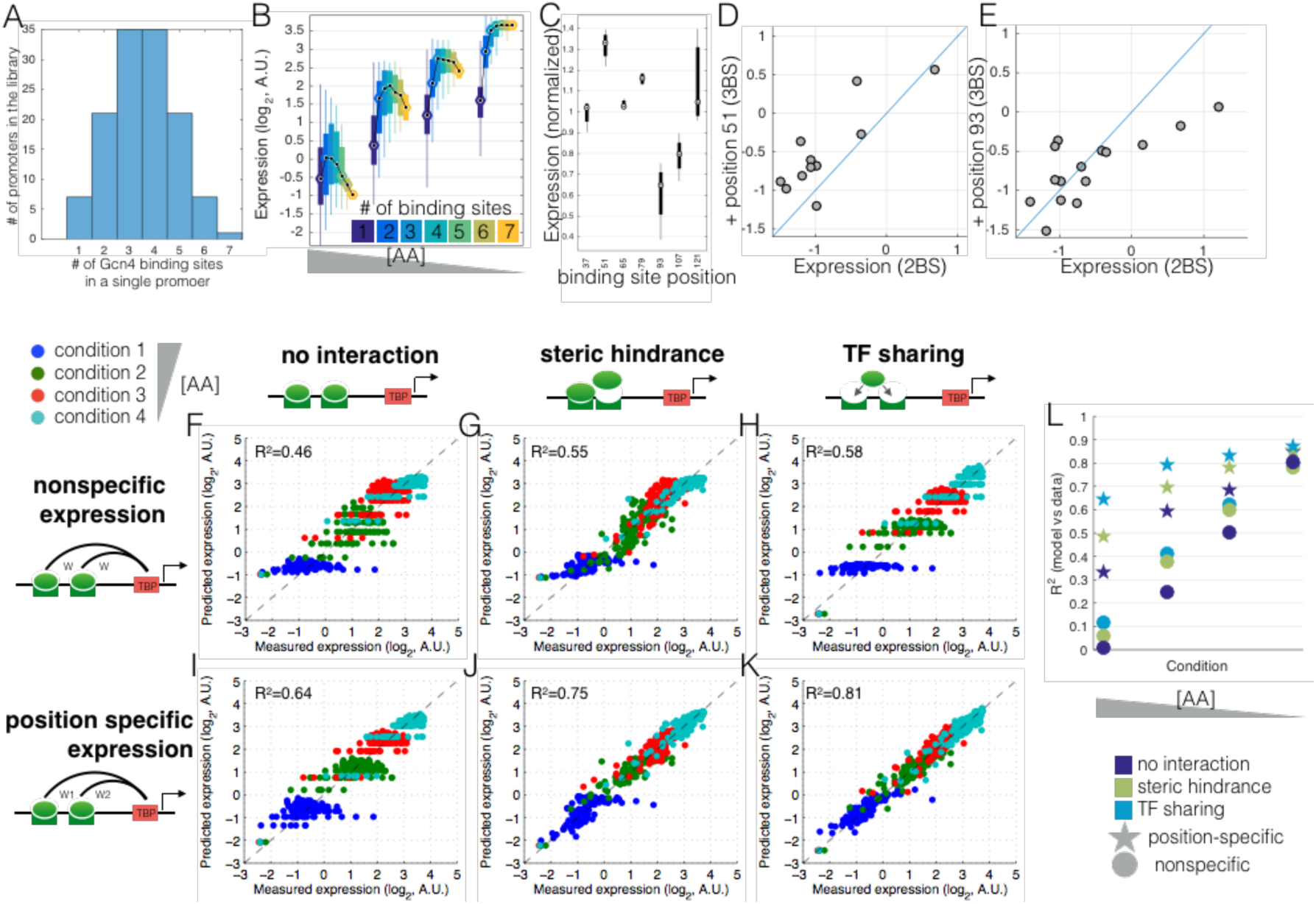
A model that incorporates TF sharing with specific position-expression can best explain expression across all amino acid concentrations. **(A)** The library contains of promoters with identical Gcn4 binding sites placed at one of seven locations in the promoter. **(B)** Shown are the measured expression levels (Y-axis) as a function of binding site number (different colors) at four AA concentrations (different groups along the X axis) for Gcn4. Each box contains data for all promoters with that number of binding sites and no other features (eg: no nucleosome disfavoring sequences or binding sites for other TFs). The black line shows the median expression level for all promoters with that number of binding sites. **(C)** Shown is the expression for each promoter with a single Gcn4 binding site, normalized so that all conditions have the same mean expression. **(D,E)** Shown is the effect of adding a third binding site (at position 51 or position 93) to a promoter that already has two binding sites. The expression of the two binding site promoter (x-axis) is graphed against the three binding site promoter (y-axis). **(F-L)** Each point shows a single promoter measured at one of four conditions (blue, green, red, cyan in decreasing [AA] order) (x-axis) and the predicted expression levels (y-axis) of that promoter, for the six different models, fitted in cross-validation to the data shown in (A), which are promoters with 1 to 7 high affinity Gcn4 binding sites (ATGACTCAT). R^2^ values were computed for absolute predicted expression on the test data. Each model includes either position specific expression (a unique weight is associated with each unique binding site position) or non-specific expression (all binding site positions share the same weight), and either no interaction, steric hindrance (a negative weight for multiple bound configurations) or TF sharing (the [TF] weight is divided by the number of sites).?

### Maximum expression is set by the amount of active TF and is limited by competition for TF molecules

If expression were a simple non-decreasing function of the number of bound TF molecules (Raveh-Sadka et al., 2009; Gertz et al., 2008), we expect expression to increase when either [TF] or number of binding sites in a promoter increases. Thus, a given expression level might be reachable by changing either one or the other, and any promoter, given enough [TF], would be able to reach a level of maximal expression set by the efficiency of transcription initiation. However, this is not what we observe in homotypic promoters. We find that the maximum reachable expression level is determined by [TF] and not by the number of binding sites (**Fig. 4A,B, Supplementary Fig. 5**). In all conditions and for all TFs, expression reaches its maximal level at 3-4 sites and then plateaus, decreases, or only slightly increases, depending on the TF, suggesting that this phenomenon is a general consequence of binding site multiplicity and not specific to a particular transcription factor.

We found that for the set of seven promoters with a single binding site placed at one of seven positions in the promoter, different binding site positions drive different levels of expression (**Fig 4C**). Furthermore, we found that when a binding site that drives high expression (eg: the site at position 51) is added to a promoter with two binding sites (generating a promoter with three sites), expression tends to increase (**Fig. 4D**). In contrast, when a site that drives low expression (eg: the site at position 93) is added to a promoter with two sites, expression tends to decrease if the expression of the two binding site promoter is already high (**Fig. 4E**).

**Figure 5.**
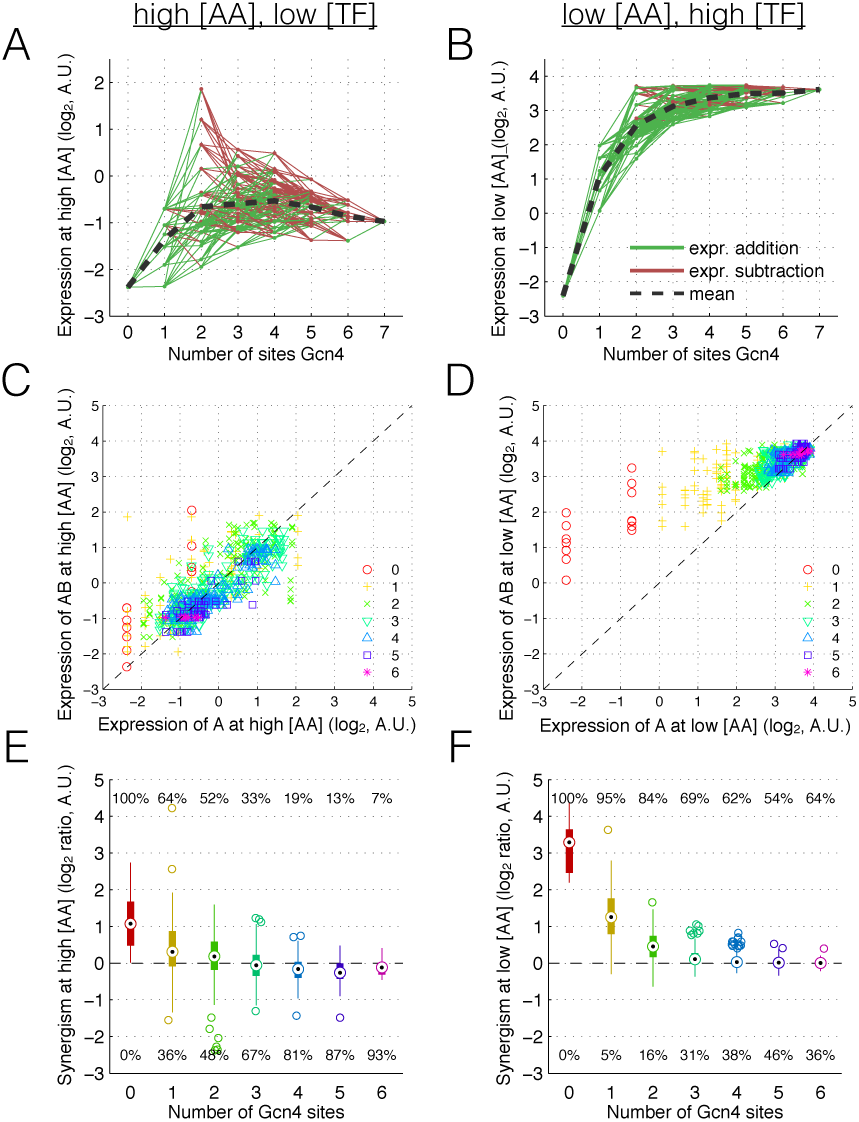
At low [TF], addition of a binding site often results in a reduction in expression. (A,B) Measured expression (and expression change, lines) when adding Gcn4 sites, for high (A) and low (B) [AA] Lines connect promoters that differ by only one binding site. Green lines indicate an expression increase when adding a binding site, red lines indicate that expression decreases. **(C,D)** A scatter plot showing the effect of adding a single Gcn4 site ‘B’ to the ‘A’ promoter, where the ‘A’ promoter has 0-6 binding sites (differently colored points) and the ‘AB’ promoter has (0-6)+1 sites. If the point is on the diagonal there is no effect of adding an additional binding site. For each AB promoter, synergism is the y-axis distance from the diagonal line. **(E,F)** Box-plots quantifying synergism — the log2 fold-change in expression when adding a binding site to promoters with 0 to 6 sites, for high (E) and low (F) [AA].

We hypothesized that the observed saturation behavior, which is most pronounced at high [AA] (low [TF]) (**Supplementary Fig. 5**), is a consequence of competition for limiting TF between binding sites that drive different levels of expression. To compare possible underlying mechanisms we used thermodynamic modeling of gene expression. In short, for each promoter, the model enumerates all possible binding configurations of TF and TBP (TATA binding protein that recruits the transcriptional machinery). The weight of each configuration is based on binding site affinities, Gcn4 concentration and interactions between bound TF molecules, and bound TBP molecules, after which the ratio between weighted TBP bound to TBP unbound configurations determines the expression (see **Methods** for details). We fitted a collection of models of increasing level of complexity to the induction curves of a set of promoters that only contain 0-7 high affinity Gcn4 binding sites. We used a 10-fold cross-validation scheme to assess each model.

A basic model (see **Methods**) in which binding to each site is independent and each site has either identical contribution to expression or with position-specific driven expression, is able to explain an increase in expression with increasing [TF] but does not fit the measured data very well (**Fig. 4F,I,L**).

We reasoned that, in order to reproduce the observed saturation, there must be negative interactions between TF binding sites within the same promoter. We examine two alternative mechanisms of binding site interaction: steric hindrance and TF sharing. The steric hindrance model accounts for a previously suggested mechanism in which a bound TF may sterically hinder the binding of a second TF molecule at a neighboring site (Struhl, 1989) by reducing the weight of configurations with multiple bound sites (Raveh-Sadka et al., 2009; Gertz et al., 2008; Giorgetti et al., 2010). The TF sharing model implements competition between neighboring binding sites by dividing the [TF] weight by the total number of binding sites. This mechanism has been observed experimentally, and results from non-specific binding and subsequent 1D sliding; two neighboring binding sites will share their TF capture area and as a consequence have the same effective binding rate as one site (Hammar et al., 2012; Mahmutovic et al., 2015).

We find that both interaction models can replicate the observed saturation effect, in which, at all [AA], adding a fourth binding site does not result in a large increase in expression (**Fig. 4**). However, quantitatively the TF sharing model better fits the experimental data.

Taken together, our results suggest that activator binding site multiplicity does not linearly contribute to expression. Our model suggests that this is due to competition between binding sites, likely due to neighboring binding sites sharing their capture area as a result of most binding events coming from 1D sliding.

### Activator binding sites can both increase and decrease expression as predicted by a model of TF molecule sharing

The above observations show that multiple binding sites contribute non-linearly to expression. In order to understand the effect of adding or removing individual activator sites in more detail we look at pairs of promoters that differ by only a single binding site. Surprisingly, in 30% of cases adding an additional Gcn4 site reduces expression (**Fig. 5**), and this effect is significantly stronger at high [AA] (55% versus 5% at low [AA], **Fig. 5E,F**), when [TF] is low. However, expression reduction is never below the minimum expression driven by the individual sites (**Supplementary Fig. 6**).

**Figure 6.**
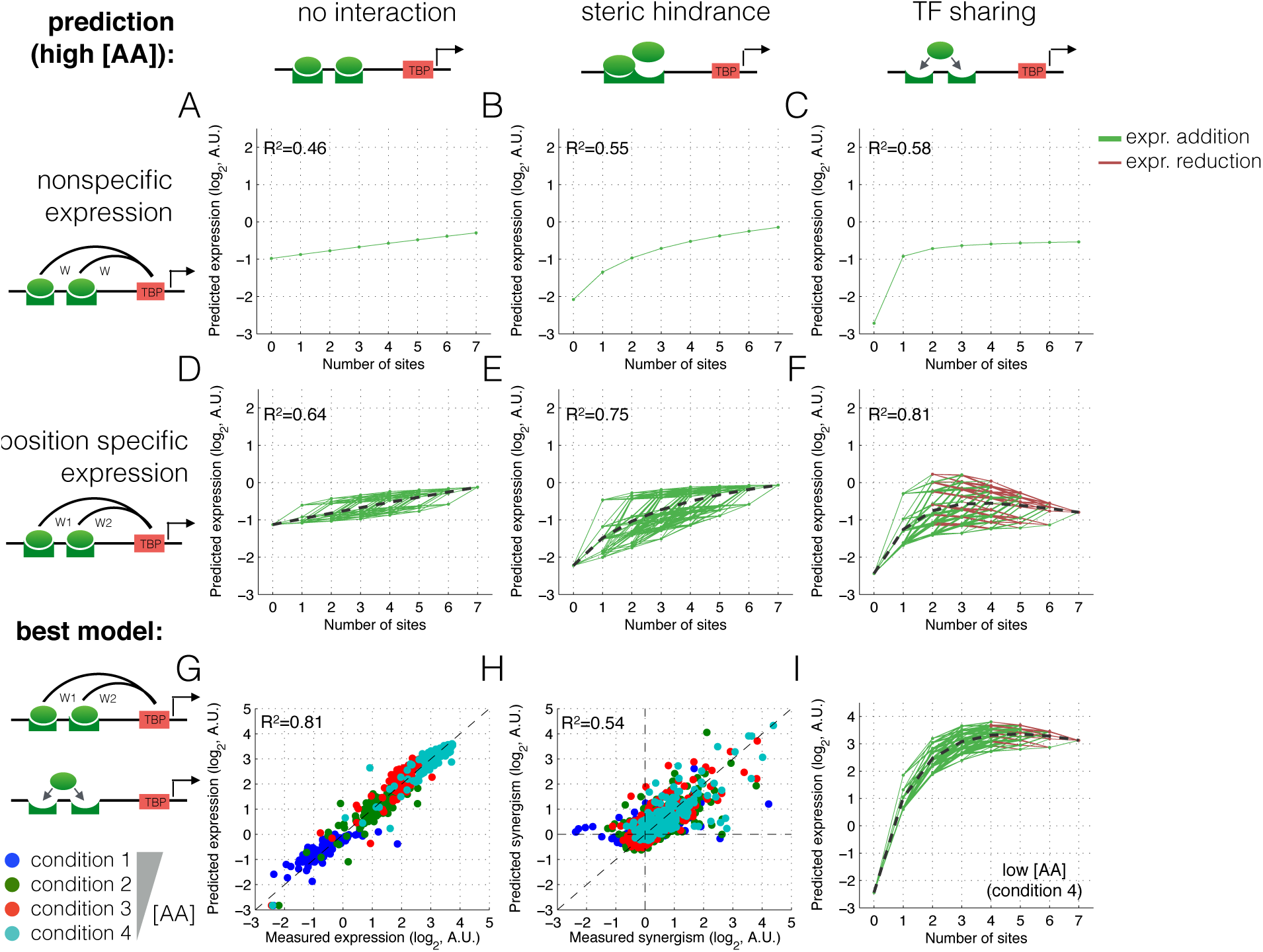
TF sharing but not steric hindrance can explain the decrease in expression due to activator binding site addition. (A-F) Predicted expression as a function of binding site number for six different thermodynamic models, fitted in crossvalidation to the Gcn4 measured data. Green lines show a predicted increase in expression upon binding site addition; red lines show a predicted decrease. R^2^ values were computed for absolute predicted expression on the test data. **(G-I)** Measured versus predicted expression and synergism for the best model at low and high [AA].

A comparison of thermodynamic models shows that both steric hindrance and TF sharing can produce expression reduction for activator binding site addition when the added site drives lower expression than the existing site. However, only the TF sharing model shows this effect at low [TF]; steric hindrance shows reduction only when [TF] is high (see **Methods**). Both steric hindrance and TF sharing models predict that the negative interaction between binding sites is stronger at closer distances. Indeed, this is the case in both expression data and in an independent measurement of the same promoter library in which TF binding to promoters was measured *in-vitro* (**Supplementary Fig. S7**). Consistent with the TF sharing model, but not with steric hindrance, this interference is strongest at low TF concentrations both *in-vivo* and *in-vitro*

The TF sharing model combined with site-specific expression best predicts absolute expression levels as well as synergism, i.e. the change in expression when adding a site (**Fig. 6, Supplementary Fig. 8–11**). In fact, the TF sharing model, given sitespecific expression, is the only model tested that can explain expression reduction at high [AA] (low [TF]).

Taken together, these results show that site addition can either increase or decrease expression. This synergism is concentration dependent. Negative synergism mostly occurs at low [TF], likely ruling out steric hindrance. TF sharing in combination with site-specific expression predicts the observed behavior: more often than not adding an activator-binding site results in a reduction of expression at low [TF].

## Discussion

In summary, we presented here a large-scale investigation of the mapping between promoter DNA sequence and dose response curves by measuring the induced gene expression of 6500 designed promoters at six growth conditions in which the regulating TFs are gradually induced.

We observe a wide range of dose-response curves in which the dynamic range is altered by changes in the affinity or number of binding sites, and expression is changed induction independently through changes in the accessibility of the promoter.

These results are confirmed by systematic mutations in either the whole promoter or only at the binding site, both affecting overall expression, but only the latter affecting the dynamic range. This suggests that random mutations (that occur more frequently outside of binding sites) are more likely to change overall expression and not the promoter’s response.

Our current and previous (Sharon et al., 2012) observation that expression saturates with increasing number of activator binding sites suggests that either TF binding or pol2 recruitment saturates. However, we observe that while expression cannot be increased by adding binding sites, expression can be increased by increasing [TF]. This argues against saturation of pol2 recruitment being the cause of the observed saturation of expression level as a function of homotypic binding sites number in each condition. We find that a model that includes competition between binding sites can quantitatively explain our observations.

We achieved further insight into the non-linear mapping between promoter configuration and dose-response by comparing pairs of promoters that differ by only a single binding site addition. This analysis revealed that at low [TF] adding an activator is more likely to reduce expression than it is to increase expression, suggesting that there is interaction between binding sites.

Expression of our synthetic Gcn4 targets maxes out at 3-4 binding sites. Interestingly, the vast majority of native Gcn4 targets have 3-4 binding sites (Schuldiner et al. 1998). The mechanistic models proposed in this paper may explain the reason for the distribution of binding site numbers in native promoters.

Our analysis of the observed dose response curves suggest that they are affected mainly by competition for TF (therefore reducing the effective local TF concentration ‘seen’ by each binding site; referred to as ‘TF sharing’) rather than steric hindrance between TF molecules. In particular, the two models behave differently with changing [TF]. While steric hindrance will have a stronger effect at high [TF] due to the increased likelihood of bound configurations, ‘TF sharing’ effects are reduced at high [TF], as the TF is no longer limiting, and this is what we observe.

To further investigate the possible mechanism that could explain the measured reduction in expression as a function of activator binding site addition, in addition to the thermodynamic model that was fit to data, we developed a toy mathematical model that describes binding site addition from 1 to 2 sites, enabling us to investigate the regimes in which addition will cause a reduction in expression (see supplementary methods). This model shows that expression reduction by steric hindrance will increase with increasing [TF], whereas reduction by TF sharing decreases with increasing [TF]. It is the latter behavior that we observe.

We note that alternative models are possible; the TF sharing model fits the data but modeling can only show that a given model is wrong, not that a given model is correct. Recently a nonequilibrium promoter-dynamics model was proposed in which TF dissociation is fast and actively driven by transcription (Coulon et al. 2013). Our results from the thermodynamic and toy models are independent of assumptions regarding dissociation. Therefore our predictions are independent of whether or not TF unbinding is a an induced non-equilibrium process. One possible alternative model that will reproduce a decrease in expression at high numbers of binding sites, specifically at low [TF], is a combination of additive activation and cooperative repression in which both the activator and repressor compete for the same binding sites. There is evidence suggesting that the transcriptional repressor Mig1 acts cooperatively (Gertz et al. 2009). While no repressors are predicted to bind with high affinity to the Gcn4 binding site (ATGACTCAT), Yap3 and Yap7 are predicted to bind weakly (de Boer et al. 2012). We hypothesized that if the Gcn4 sites have repressive potential, then site addition can cause expression reduction below the level driven by the other sites. We find that, while addition of a binding site often results in expression below the maximum of the expression driven by the individual sites this expression is always greater than the minimum expression driven by the individual sites. The added site can, at most, reduce expression by an amount that the other sites drive, and can never repress beyond that level. In other words, we find that to remove expression first expression has to be added. This is a strong prediction of the TF sharing model and is not predicted by the cooperative repression model.

A second model that can explain the observation that expression reaches a maximum at around three binding sites is that having more than three bound TF molecules does not increase recruitment of RNA polymerase. It is likely that beyond some number, additional bound transcriptional activators do not contribute to increased expression at a single promoter. However, this model cannot qualitatively explain our observation that, for all four TFs, the expression from 3 binding sites is lower at higher [AA] (lower [TF]). The ‘activator saturation’ model predcts that the same maximal expression could be reached at all amino acid concentrations, but that it might require more binding sites at lower [AA]. This is not what we observe. Moreover, while on average expression saturates at three sites, this is not always the case. Figure 5A shows that going from three to four sites can both increase (green lines) and decrease (red lines) expression; expression rarely remains constant, likely ruling out the ‘activator saturation’ model.

Taken together, we have found a strong non-linear mapping between promoter architecture and dose-response, that, by assuming competition between binding sites, we are able to accurately predict from DNA sequence alone. In specific, our model points to a reduction in effective local [TF] (per binding site) due to overlapping capture areas. When [TF] is limiting the effective search time (the time it takes for a TF to find its binding site) is not significantly reduced when another site is added close to an existing one, since search time is dominated by the total capture area. In the regime where [TF] is high more sites bind more TFs and thus have the ability to drive higher expression.

Our model is also consistent with recent *in-vitro* results performed using the same set of promoters showing that at low [TF], multiple Gcn4 binding sites increase the likelihood of TF binding, but do not increase the number of TF molecules bound to a single molecule of promoter, while at high [TF], adding more binding sites does increase the number of bound molecules (Levo et al., 2015).

Competition for limiting TF, in both one and three dimensional space, may also explain some previously unexplainable results regarding titration by large arrays of extraneous TF binding sites. Lee & Maheshri (Lee and Maheshri 2012) found that contiguous arrays of tetO binding sites bind less TF than do non-contiguous arrays, and that contiguous arrays are less efficient at titrating away TF. Our reanalysis of their data shows that this effect is strongest at low [TF], suggesting that TF sharing may be occurring at these arrays as well, either in 1D space or in 3D space. Splitting the array of decoy binding sites in half results in a larger decrease in expression at low [TF] than at high [TF] (**Supplementary Fig. 12**), as expected from a model in which large number of binding sites spread throughout the genome (at the promoter of interested and at the decoy sites) are sharing a limiting number of TF molecules.

The yeast genome, which has densely packed genes for a eukaryote, has several promoters (e.g.: Cln3) that are longer than 1kb. Yet 20 TF binding sites could, in theory, be packed into less than 200bp. Intriguingly, the GAL genes, which are highly induced by a large and rapid increase in the active amount of Gal4, tend to have only one or two nucleotides between the sites. In contrast, genes activated at the G1->S transition (e.g.: Cln2) have TF binding sites that are spaced further apart. It was recently shown that Cln3, the protein that activates the TFs bound to the Cln2 promoter is present in limiting concentrations(Wang et al., 2009); the spacing between binding sites may reduce the effect of sharing. Binding site spacing is known to be influenced by physical interactions between TFs (Kazemian et al., 2013). Here we suggest that TF sharing between closely spaced binding sites is an additional force acting upon the evolution of promoters. Binding sites for some TFs, especially those with long 1D sliding ranges(Gorman and Greene, 2008; Slutsky and Mirny, 2004) may need space for maximal TF occupancy at low [TF]. Dense clusters can be used to create a highly responsive behavior (large dynamic range) and less dense clusters might create overall high expression also at low [TF]. Our results suggest that TF sharing can play an important role in determining the response of a promoter to changes in [TF], and therefore influence the evolution of binding site configurations.

## Methods

### Promoter library design construction and measurements

We used a previously described library of 6500 bar-coded sequences, each with a specific combination of TF binding sites and poly-A tracts(Sharon et al., 2012). These sequences were cloned into a pKT103-derived plasmid upstream of a yellow fluorescent protein (YFP) and transformed into the yeast strain Y8205(Tong and Boone, 2006). We then grew the pooled library in six different growth media each with a different concentration of amino acids: synthetic minimal media with glucose and the amino acids His & Leu with a 2^11^, 2^6^, 2^4^, 2^3^, 2^2^ or 2^0^ fold dilution of amino acids. Cells from each growth condition were sorted by FACS according to expression (YFP / mCherry) into 16 bins. DNA from condition and bin was PCR amplified using unique primers so that each resulting sequencing read has three bar codes: promoter, condition, expression bin. The distribution of reads across the bins for each promoter and condition enables us to derive an expression level as previously described(Sharon et al., 2014). Thus, we achieve, for each promoter expression across the conditions.

A Gcn4-GFP ura3::TEFpr-mCherry strain was grown overnight in SCD-HL, resuspended in SCD-His-Leu or SCD, and then the SCD was serial diluated into SCD-HL, resulting in different concentrations of His and Leu. GFP and mCherry were measured using a BD Fortessa flow cytometer using FITC and PE_TexasRed filter sets.

### Expression normalization

We observed condition specific expression differences that did not appear to stem from biological differences. For example, even the promoters that were not induced (such as Gal4 targets) varied, though slightly, across conditions in a non-monotonic manner. These differences likely stem from day-to-day and experimental variability, as each condition was a separate batch and was sorted on different days. To correct for this effect we subtracted from all promoters the median expression of all Gal4 targets, thus removing this technical variability. All analysis was carried out on the normalized expression values.

### Growth conditions

Because the two lowest and two highest [AA] conditions induce similar expression we combined them to get a more robust expression measurement. Thus, for the analyses in which we compare low to high [AA] we use the average of the two lowest and the average of the two highest [AA] conditions. In the analyses in which we compare four conditions we use the previous two plus the middle two [AA] conditions.

## Acknowledgments

This work was supported by grants from the European Research Council and the US National Institutes of Health to E. Segal and a grant from the Agència de Gestió d'Ajuts Universitaris i de Recerca (AGAUR) to L. Carey. D. van Dijk was supported by NWO Rubicon fellowship 825.14.016.

## Figures

**Supplementary Figure 1.**
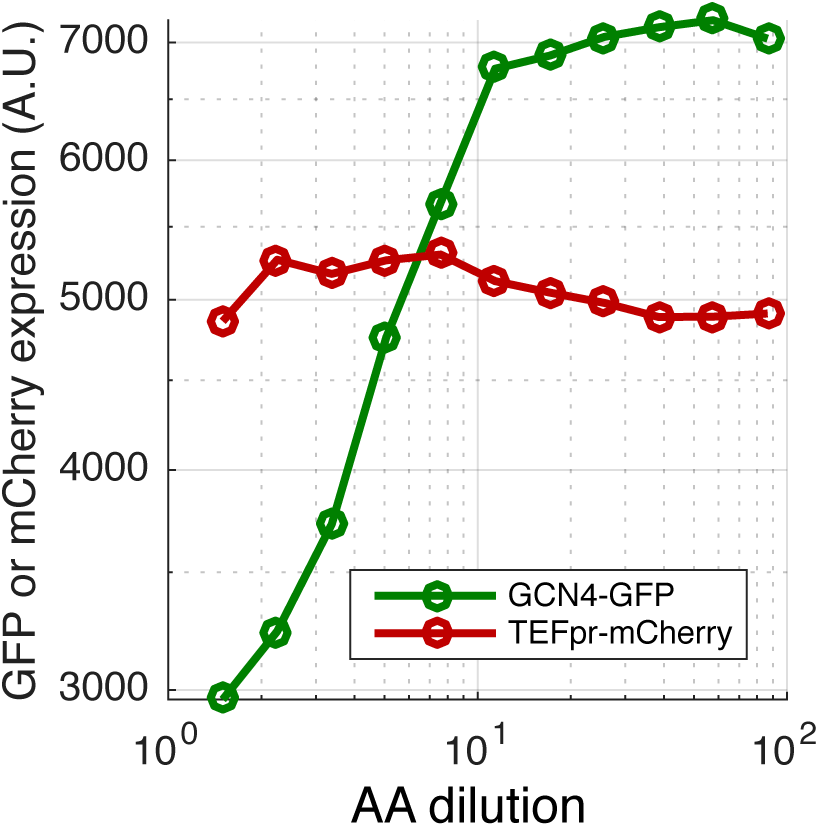
Amino acid starvation induces Gcn4 transcription factor levels. Shown is a serial dilution of amino acid concentration and the levels of Gcn4-GFP and a control TEF promoter driving mCherry. The Gcn4-GFP-HIS3 *ura3*::TEFpr-mCherry-URA3 strain is from the GFP collection. Cells were grown in SCD, washed, diluted into varying concentrations of SCD-HL media, and grown for eight hours.

**Supplementary Figure 2.**
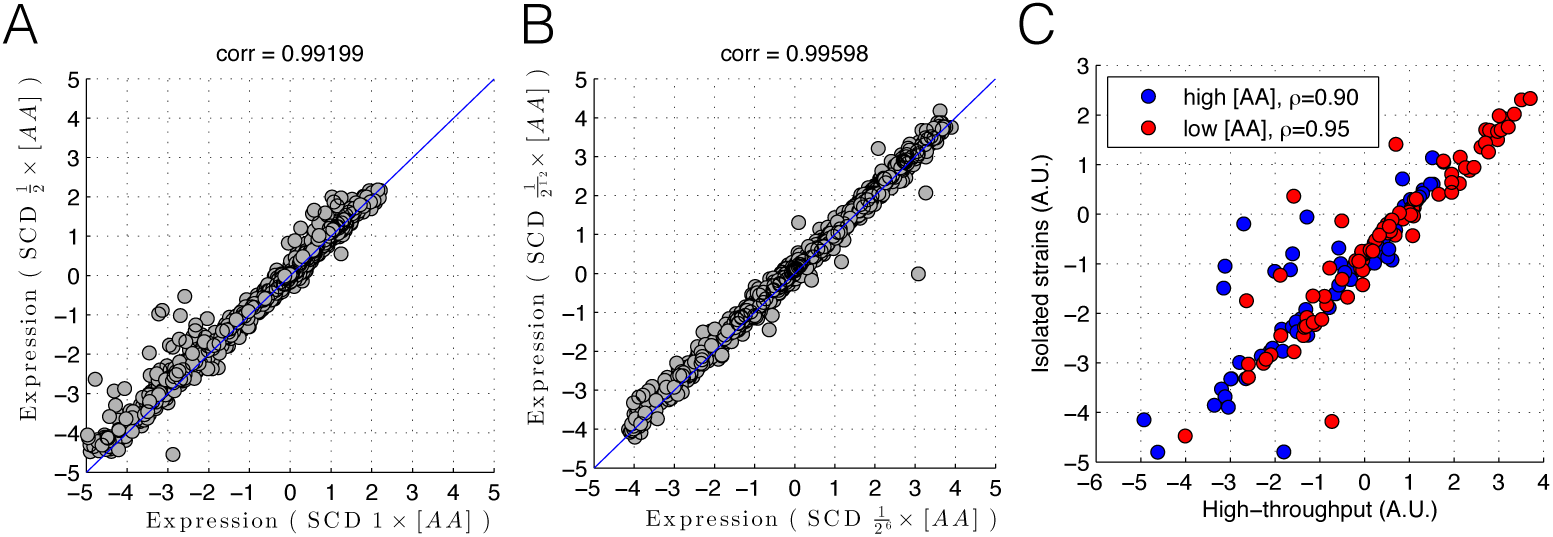
Reproducibility between biological replicates and between isolated and high-throughput measurements. Shown are biological replicates of the high-throughout measurements (A, B) and isolated strains measured in plate reader (C). (A) shows condition 1 versus 2 and (B) shows condition 5 versus 6. These conditions were measured on different days. (C) shows 96 promoters that were isolated form the pooled library and measured using a plate-reader in high and low [AA] (equivalent to conditions 1 and 6). The X-axis shows expression measured in the high-throughput pooled experiment and the Y-axis shows the expression measured using the plate-reader for individual strains.

**Supplementary Figure 3.**
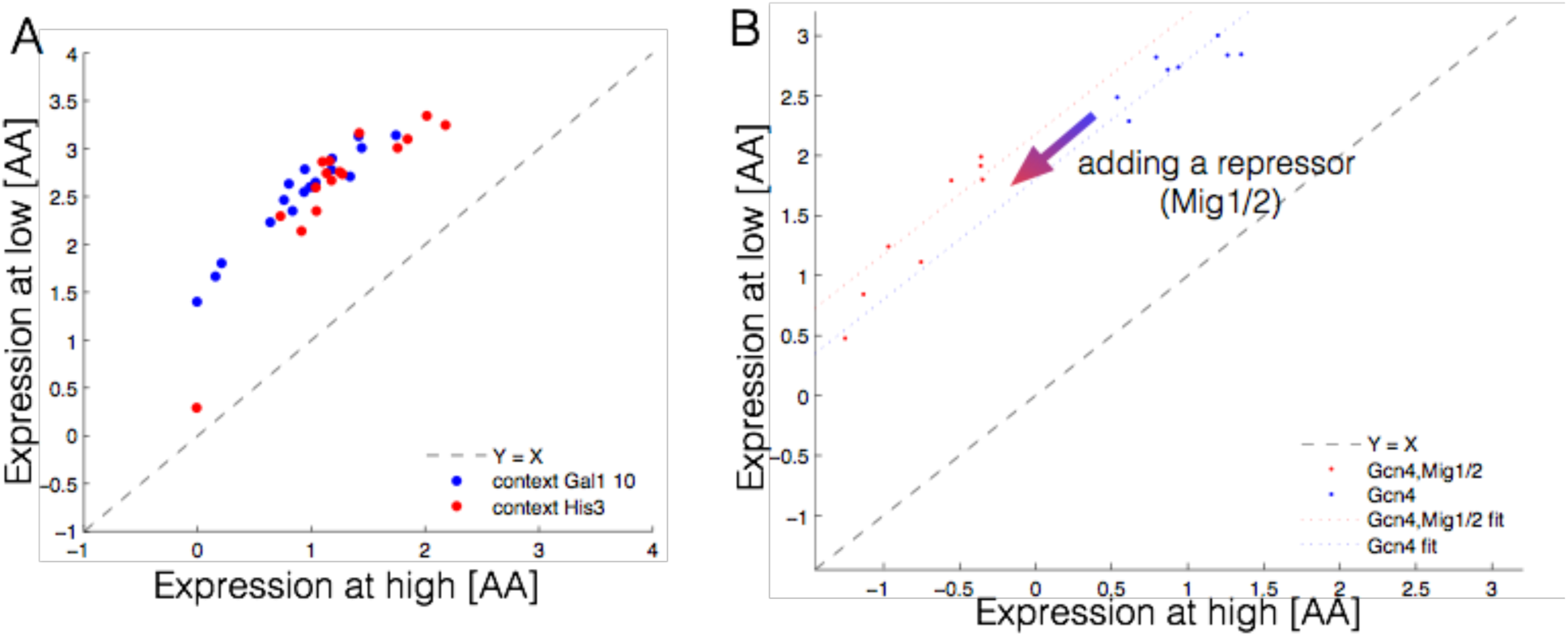
Changing the promoter sequence context or addition of a binding site for a repressor changes expression without changing dynamic range. **(A)** Shown are expression at high and low [AA] for a set of promoters with a single Gcn4 binding site placed at different locations with the HIS3 (red, low nucleosome occupancy) or GAL1,10 (blue, high nucleosome occupancy) promoter context. (B) Shown are expression at high and low [AA] for a set of promoters with a single Gcn4 binding site placed at different locations (blue) and the same promoters with a Mig1/2 repressor-binding site added to the −36 position in the promoter. Addition of a repressor binding site moves expression along the line X=Y, and therefore affects [TF] independent expression.

**Supplementary Figure 4.**
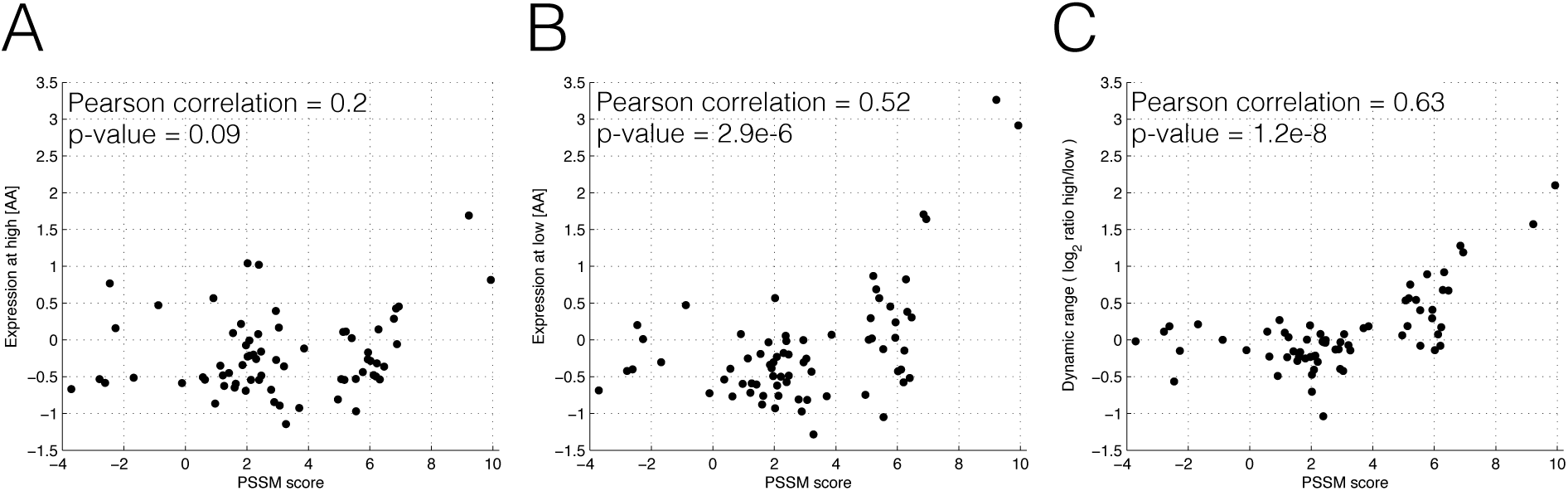
Mutations in the Gcn4 binding site that decrease the PSSM score decrease [TF] dependent expression. **(A)** Expression at high [AA] for a set of promoters that differ only by the sequence at the single Gcn4 binding site in the promoter. **(B)** Expression at low [AA] for a set of promoters that differ only by the sequence at the single Gcn4 binding site in the promoter. **(C)** Dynamic range for each of the promoter calculated from the data shown in (A) and (B).

**Supplementary Figure 5.**
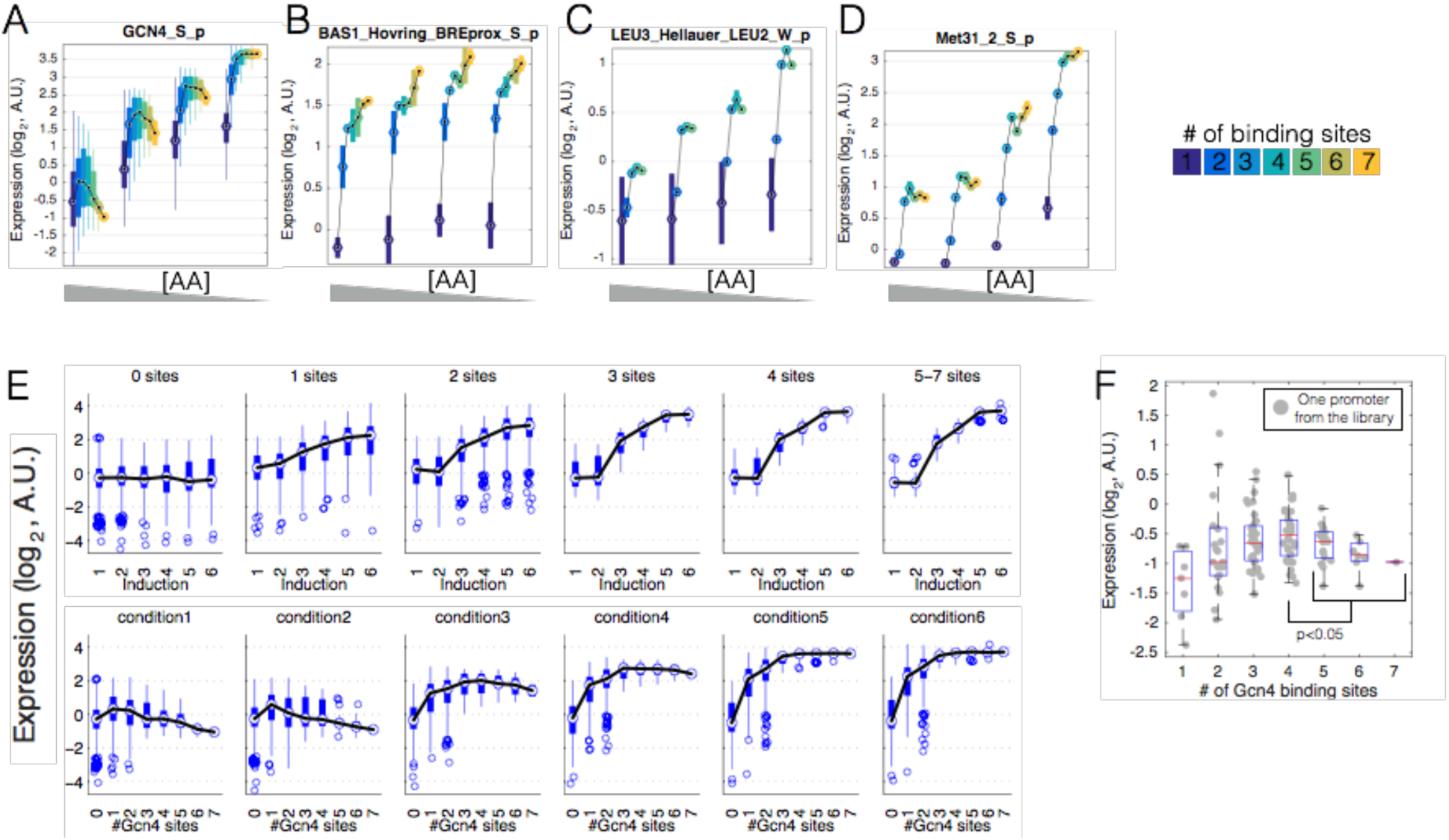
Expression saturates with binding site number for multiple transcription factors. (A-D) Shown are the measured expression levels (Y-axis) as a function of binding site number at four AA concentrations (see Methods) for Gcn4, Bas1, Leu3 and Met31 TFs. Each box is the set of promoters with 0 to 7 binding sites (different colors) measured at one of four conditions (x-axis). Black lines show the median expression per condition along the number of binding sites. **(E)** Various ways of graphing the Gcn4 binding site data. **(F)** Expression decreases when changing from 4 to >4 binding sites.

**Supplementary Figure 6.**
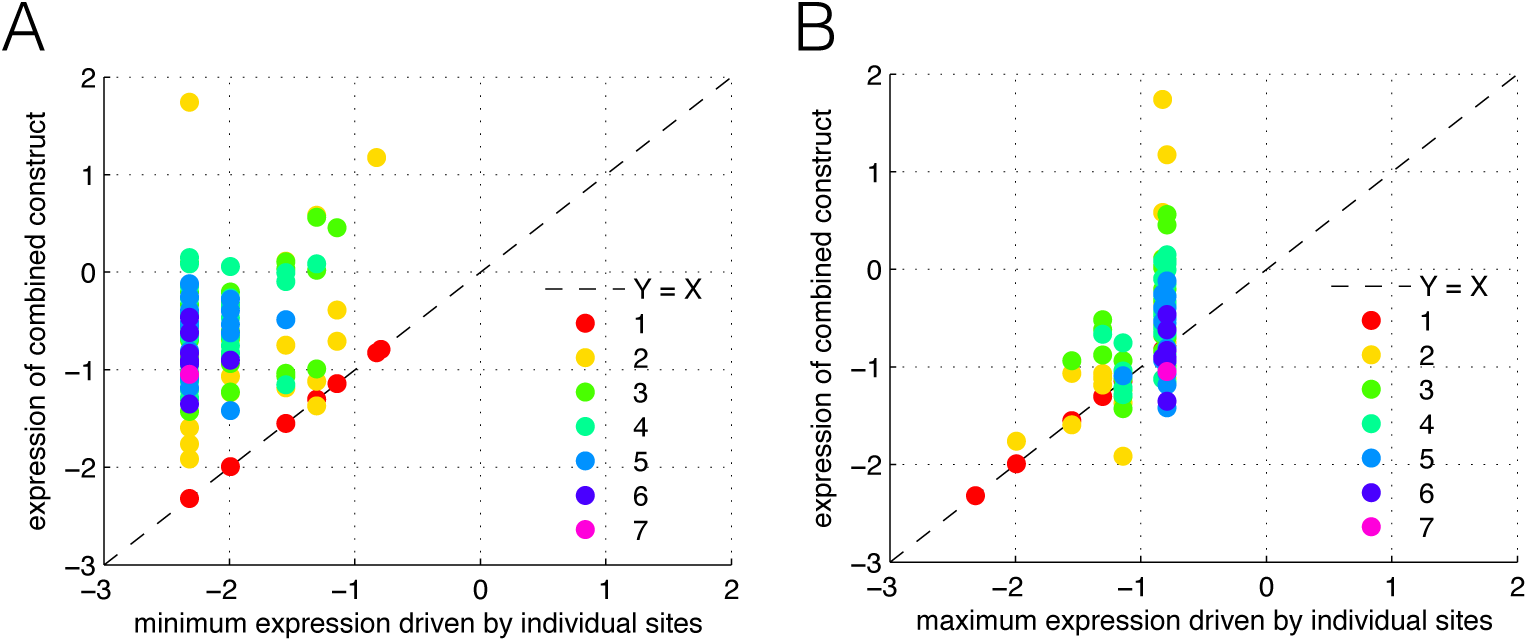
Repression is never below the minimum expression driven by the individual sites. Shown is the minimum (A) and maximum (B) expression of the individual sites (X-axis) versus the expression of the promoter in which the sites are combined (Y-axis). Thus, each point is a promoter with N sites (1-7, colors) for which the expression is shown on the Y-axis. On the X-axis the minimum or maximum expression is shown computed on the expression values of the set of promoters that have only one binding site, namely the same as is present in the promoter that is depicted, e.g. for a promoter with 3 sites S1, S2 and S3 the minimum or maximum is computed on the expression values of the promoters that have only site S1, S2 or S3.

**Supplementary Figure 7.**
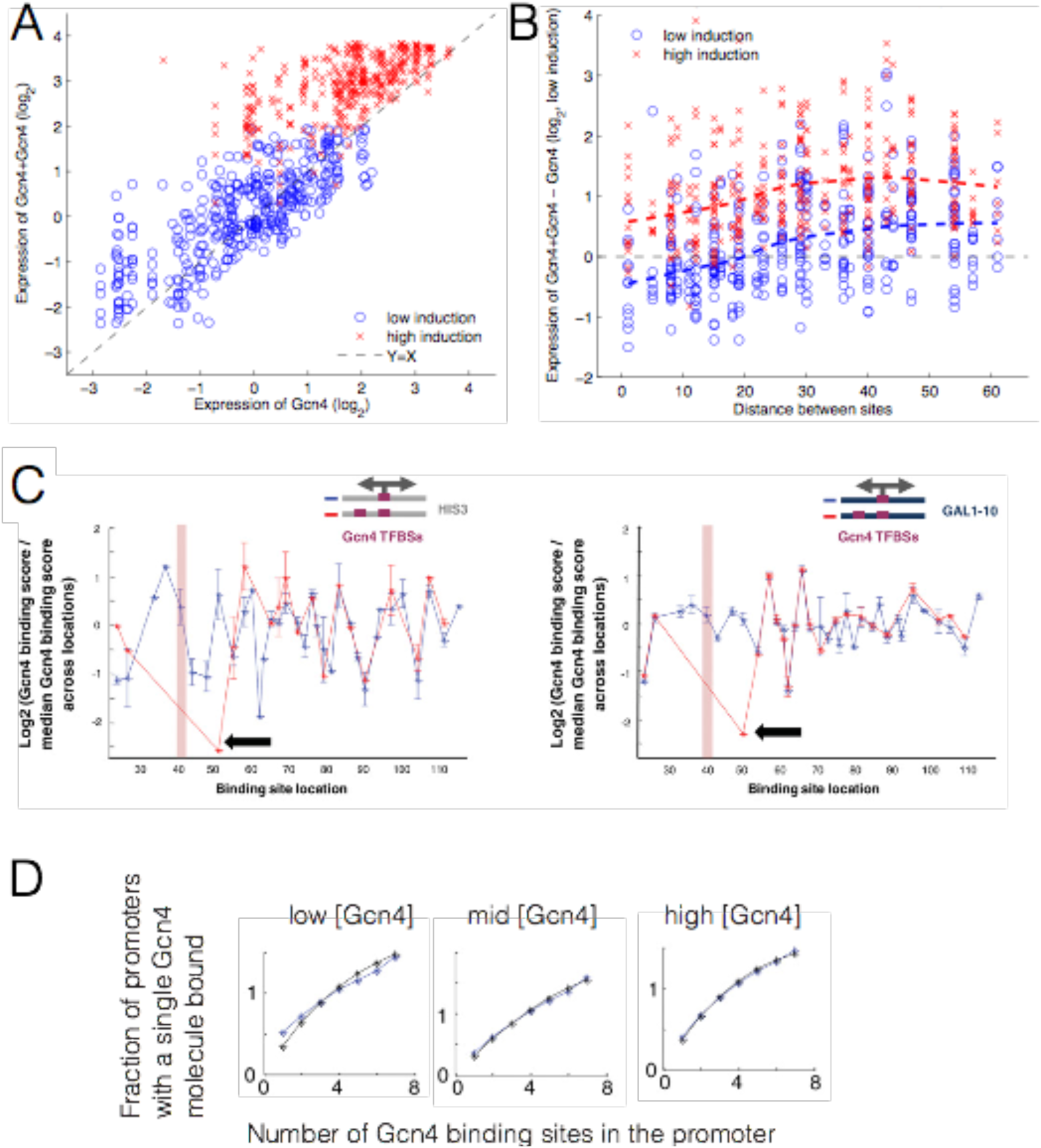
In-vivoo expression and in-vitroo binding increase less than expected when adding a second near-by binding site. **(A)** Expression change as a function of adding a second Gcn4 site. Each point shows expression for a promoter with one Gcn4 site (X-axis) and the same promoter with a second site added (Y-axis). **(B)** Expression change for each promoter pair is shown (Y-axis) as a function of the distance between the two sites (X-axis). Dashed lines show the median change in expression across all binding site additions. **(C) Levo et al. 2015. Figure 3.** A set of sequences with a strong Gcn4 site placed at different locations along a specific sequence context, either in the presence of an additional strong Gcn4 site located in a fixed location (with the pink rectangle marking the location of the center of this site) (in red) or without this additional site (in blue). Shown is the log2 of the ratio of the binding score attained by each sequence (with the x-coordinate marking the location of the center of the site) divided by the median binding score across all sequences in this set. The black arrow points to a sequence where the 9-bp sites are separated by a single bp. Sequences with Gcn4 TFBSs of 9 bp placed along the HIS3-derived context (left panel) and along the GAL1-10-derived context (right panel). **(D) Levo et al. 2015. Figure 3.** For a set of sequences with all possible combinations of one to seven binding sites for Gcn4 in seven predefined locations, the average frequency of sequences with a single molecule of bound Gcn4 is shown as a function of the number of sites within the sequence (in blue). The predictions of these dependencies are based on a simple thermodynamic model assuming multiple TF binding events are independent and are also plotted (in black). At low [TF] (left panel), the amount of bound Gcn4 increases less slowly than is predicted from a thermodynamic model lacking negative interactions between TF binding sites.

**Supplementary Figure 8.**
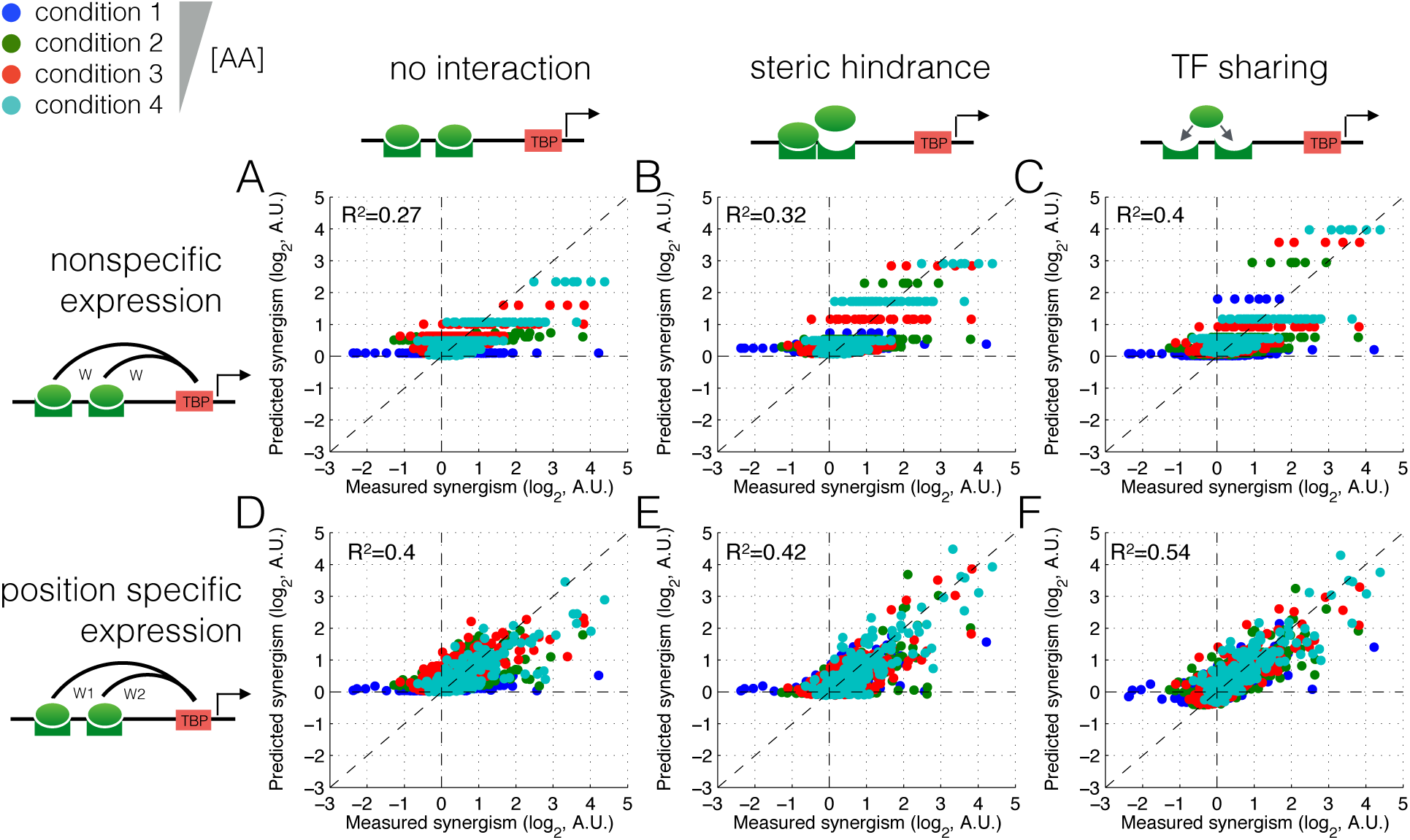
TF sharing but not steric hindrance can best explain synergism at all amino acid concentrations. (A-F) Measured versus predicted synergism for each [AA] (different colors) for six different thermodynamic models, fitted in 10-fold cross-validation to the measured data. Note that TF sharing is the only model that can generate negative synergism.

**Supplementary Figure 9.**
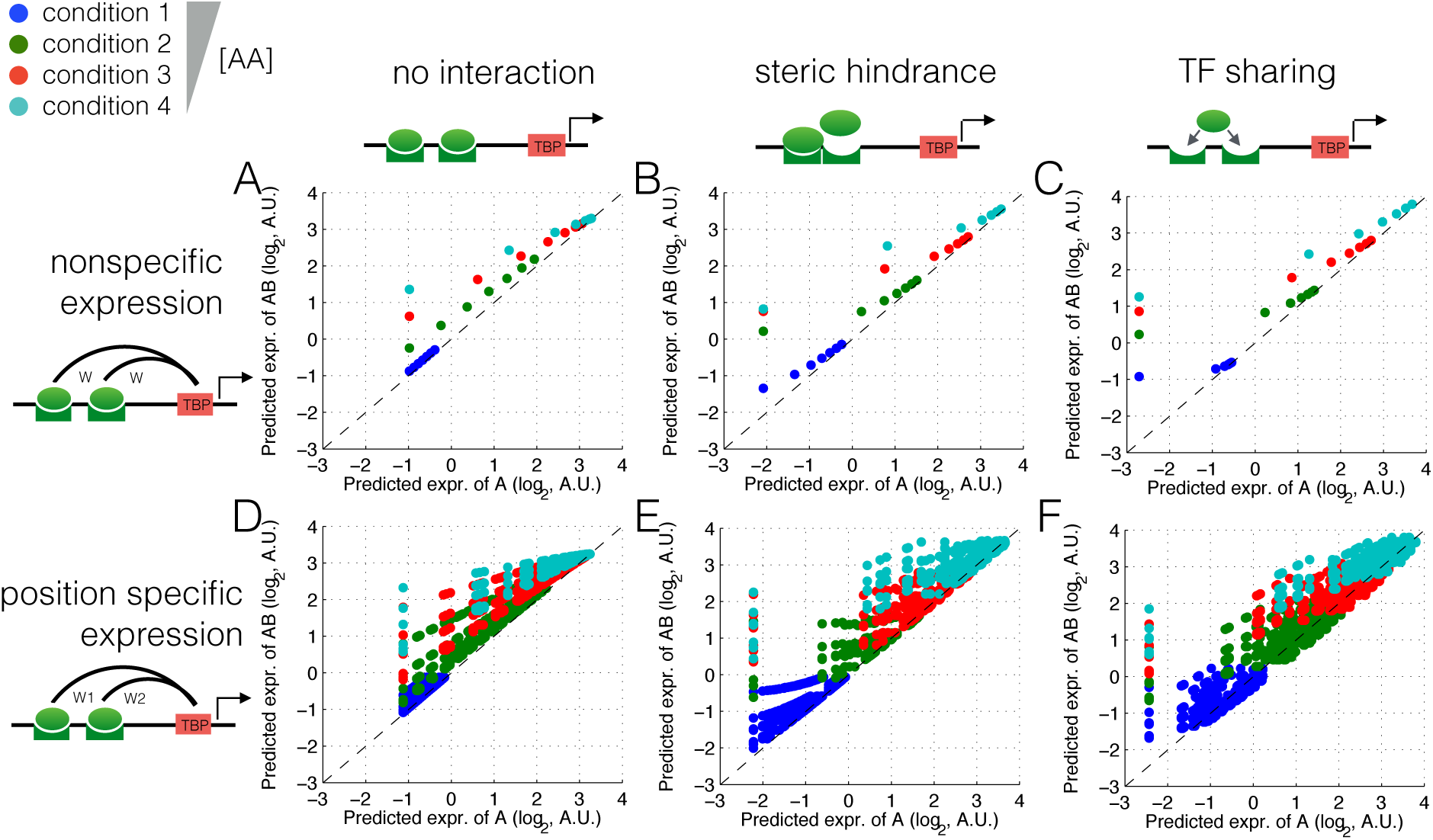
Only the TF sharing model is capable of producing strong negative synergism. Shown is predicted expression of a single binding site (A) versus expression following the addition of another binding site (AB), for all six thermodynamic models. The top row of models have fewer points visible because they don’t have position-specific expression, so all promoters that differ only by the position of binding sites will have the same expression.

**Supplementary Figure 10.**
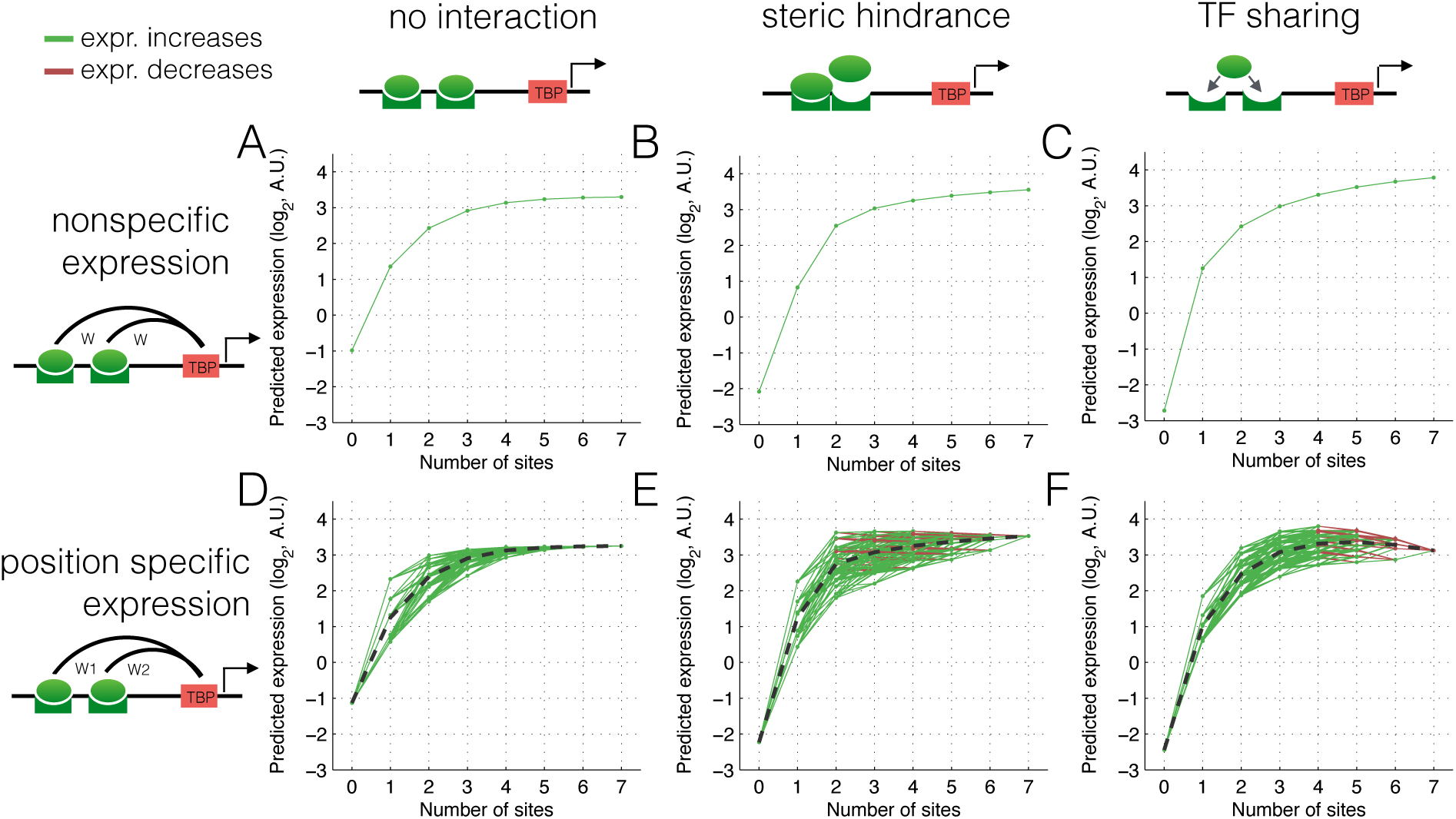
TF sharing and steric hindrance can explain expression saturation and small expression reductions at low [AA]. (A-F) Predicted expression at low [AA] is shown as a function of binding site number for six different thermodynamic models, fitted in cross-validation to the Gcn4 measured data. Green lines show a predicted increase in expression upon binding site addition; red lines show a predicted decrease.

**Supplementary Figure 11.**
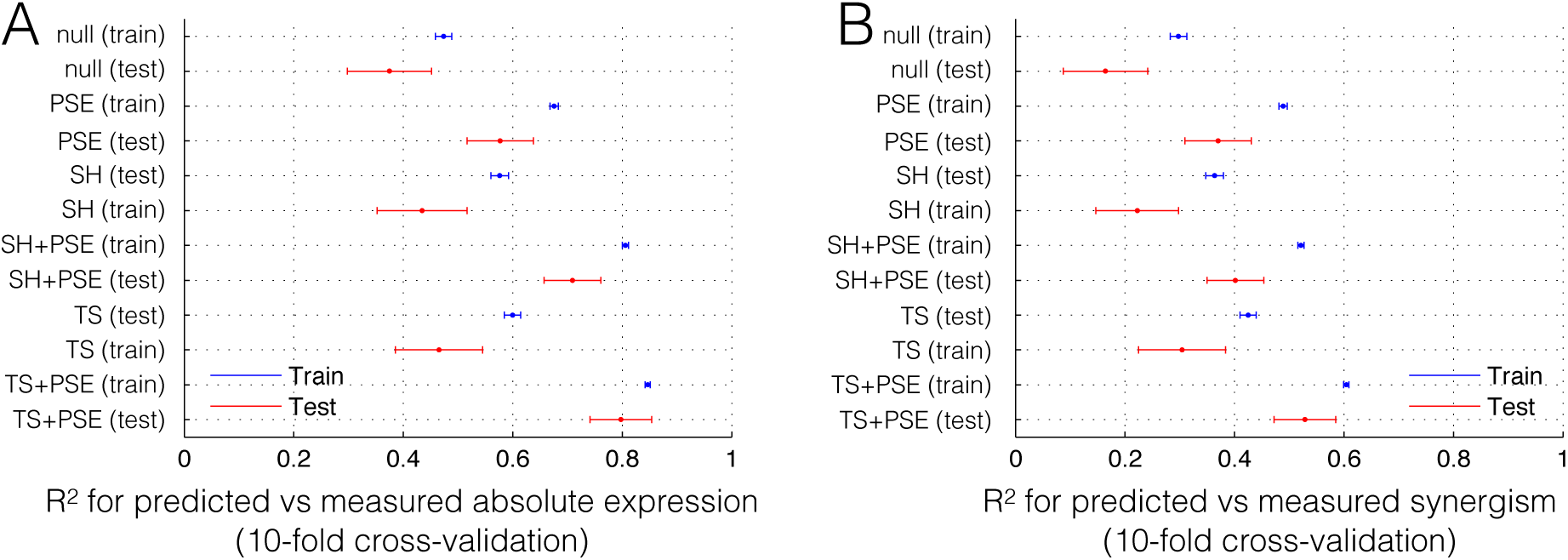
The cross-validated thermodynamic models do not exhibit excessive over-fitting. All six thermodynamic models were fit to absolute expression level at all induction points. Shown is the R^2^ for predicted versus measured expression level **(A)** and synergism **(B)** for each training (blue) and test (red) data set.

**Supplementary Figure 12.**
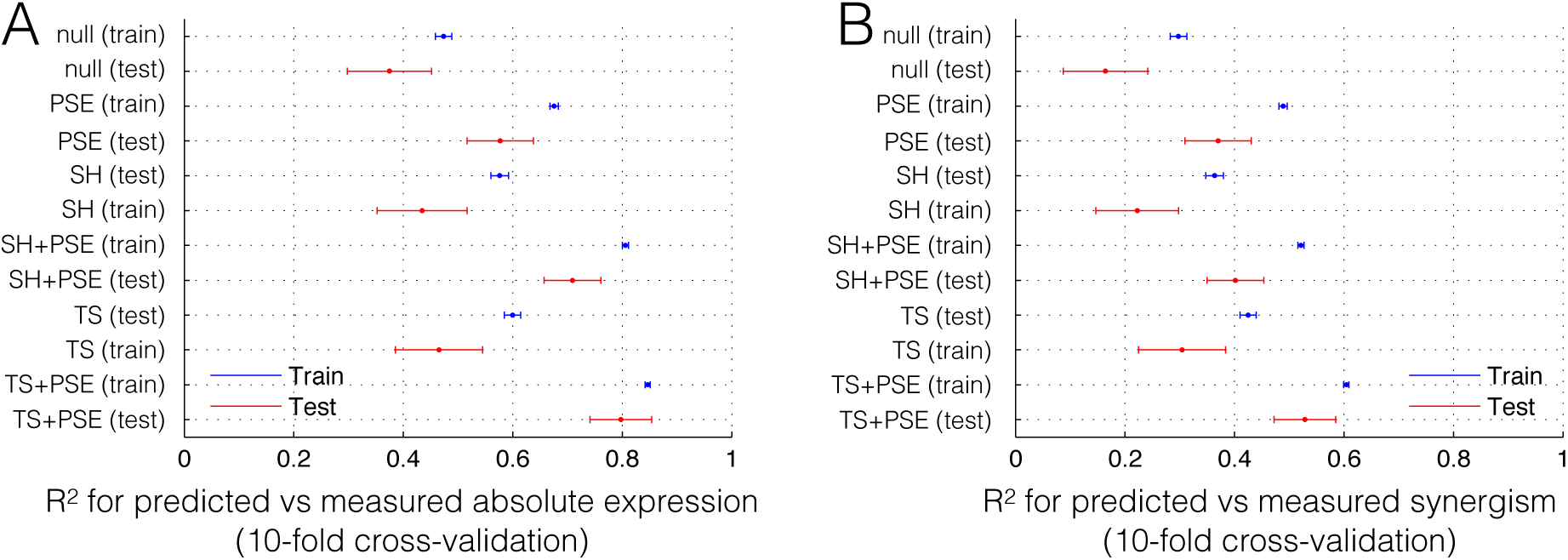
Addition of large arrays of excess binding sites titrate away limiting TF molecules independent of the number of binding sites in the array. Lee & Mahreshi used a strain in which YFP is activated by the tTA transcription factor, which is inhibited by doxycycline. Addition of doxycycline reduces the effective TF concentration. Into this strain they integrated into the genome either one or two arrays of extraneous tTA binding sites, with the arrays having 67, 127 or 240 binding sites. Data from Figure 3 were replotted to compare strains with approximately equal numbers of binding sites **(A,B)** or equal numbers of loci containing extraneous sites **(C,D)**. Splitting the binding sites across two loci increases the distance between the binding sites and results in a large decrease in expression (increase in titration of TF). However, adding extra binding sites to an existing locus results in little or no decrease in expression (no increase in TF titration).

## Supplement

### Thermodynamic model fitted to data

Our aim is to predict gene expression from promoter DNA sequence features across several transcriptional-activator concentrations. In specific, our input sequences are 128 promoters that contain 0 to 7 activator Gcn4 binding sites in 7 predefined positions for all (2^7^) possible configurations. We use a thermodynamic model (Shea & Ackers 1985, Bucheler Hwa 2003, Gertz Cohen 2009) that predicts gene expression by enumerating all promoter configurations and computing the probability of TBP binding (*P(TBP)*), which then is converted to gene expression via a sigmoid function (Eq.1).

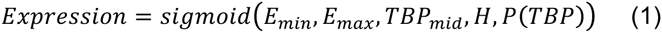

Where *E*_*min*_ and *E*_*max*_ are the minimum and maximum expression respectively, *TBP*_*mid*_ is the *P*(*TBP*) at which expression is half maximal and *H* is the hill-coefficient and describes how switch-like expression is as a function of TBP binding. This sigmoid function describes both transcription and translation.

Both expression (as a function of P(TBP)) and [TF] (as a function of growth condition) are modeled with the same sigmoid function (Eq.2).

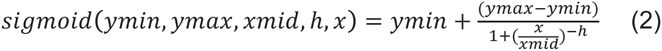

The probability of TBP binding is computed through the relative weight of TBP-bound to TBP-unbound configurations (Eq.3).

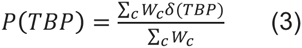

Where *W*_*c*_ is the statistical weight of configuration *c*, which is computed as (Eq.4):

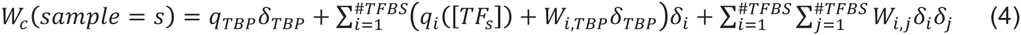

Where *δ*_*i*_ is an indicator function of whether the TF is bound in site *i*, *W*_*i,j*_ reflects an interaction between bound TF molecules and would be positive for cooperative interaction. In our case we use this as a negative weight to model steric hindrance. *W*_*i*,*TBP*_ is the interaction weight of bound TF to bound TBP and is either specific to each site position or shared across all positions. This weight can be interpreted as the contribution of a bound TF to initiating transcription.

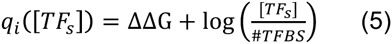

ΔΔG is the binding affinity per position. The change of Gibbs free energy is similar between sites (same sequence motif) thus we assume it to be constant across positions.

When we assume that sites act independently or for steric hindrance #TFBS = 1. When we assume TF sharing we set #TFBS to the number of binding sites per promoter.

Finally we assume that the TF concentration follows a sigmoid as a function of the conditions (Eq.6).

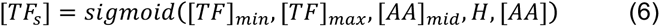

Where [*AA*] is the amino acid concentration that is varied across conditions (see Methods). ^[*TF*]_*max*_^ and ^[*TF*]_*min*_^ are the maximal and half-maximal TF concentrations. *H* is the hill-coefficient.

For the steric hindrance model we use a negative weight for TF-TF interactions. Thus, W_*i,j*_ (Eq.4) becomes W_coop_ (see model parameters below). For the TF sharing model we compute an effective TF concentration ([TF]_eff_) by dividing the [TF] by number of binding sites. The amount of sharing is set by W_coop_ as follows: [TF] _eff_ = [TF] / (N_tf_ * W_coop_), where N_tf_ is the number of binding sites.

### Model parameters

Next we describe the free parameters of the model that were fitted in cross-validation. Accompanying data files (*_params.tab) contain the fitted values for each model.

*Binding affinities:*

Q_GCN4: binding affinity of Gcn4 (for sequence motif TGACTCA)
Q_TBP: binding affinity of TBP
*TF concentration:*

C_max: Maximum [TF]
C_mid: [AA] for which [TF] is half-maximal
C_H: Hill-coefficient for [TF] as a sigmoid function of [AA]
C_min: the minimum [TF] which is fixed to 1, since this parameter is redundant with the binding affinity of Gcn4.
*Interaction weights:*

W_GCN4_TBP: Interaction weight of bound Gcn4 with bound TBP
For position specific expression this is modeled with 7 separate parameters:
W_GCN4_TBP_PosX, where X = {22,36,50,64,78,92,106}, which is the position relative to the ATG.
*Expression from TBP occupancy:*

TBP2Exp_min: minimum expression TBP2Exp_max: maximum expression TB2Exp_mid: (P)TBP for which expression is half-maximal
TBP2Exp_h: Hill-coefficient
*Steric-hindrance:*

W_coop: (Wtf-tf) TF-TF interaction weight
*TF sharing:*

W_coop: (Wtf-tf) weight that determines to what extent [TF] is shared between neighboring sites.

### Toy model of activator site addition

Both steric hindrance and TF sharing may explain, in certain regimes, reduction of expression from adding an additional binding site. To better understand how and when these models that include interactions between binding sites predict expression reduction, we developed a toy thermodynamic mathematical model that describes expression change as a result from binding site addition. We model the expression change from one to two activator binding sites by enumerating all possible binding configurations of the TFs and compute expression from the configurations that have at least one TF bound. Each of the two possible binding sites has its unique contribution to expression (i.e. interaction with the transcriptional machinery). The weight of each configuration is computed from the weight of the TFs (affinity * concentration) (parameter *Wtf*). Per configuration expression is the sum of the product of TF weights and site-specific expression. Total promoter expression is then computed from the sum of expression over all configurations. The unbound configuration has weight 1 (arbitrary constant). We then assume (1) expression driven by each site is independent, or (2) steric hindrance: the double bound configuration has weight 0, or (3) TF sharing, in which, for the promoter with two sites, [TF] = [TF]/2. Next we compute expression change from site addition by subtracting the expression from two sites from the single site promoter. To determine which parameter regimes enable expression to decrease, we solve this equation and get the boundary condition in which site addition does not change expression. We can then easily find the regimes in which expression increases or decreases, namely when the expression driven by the second site (E2) is higher or lower, respectively, than the boundary condition.

Thus, the expression change as a result of addition of a second site, for independent expression is:

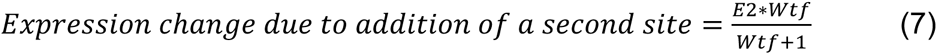

This value is always positive, thus ruling out independence. For steric hindrance,

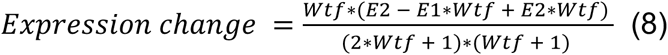

Solving gives us the boundary condition 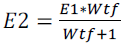 for which expression does not change. At low [TF] (low Wtf) this limit is low, thus at low [TF] we can only reduce expression when E2 << E1. This limit goes to E2 = E1 when [TF] goes to infinity, thus at high [TF] it becomes easier to reduce expression, i.e. only a small difference between E1 and E2 is necessary. So, even at high [TF] reduction will occur when the added site drives lower expression. This is not what we observe. Experimentally, the frequency of reduction goes decreases at high [TF], steric hindrance predicts the opposite, ruling out steric hindrance.

Finally for TF sharing we get

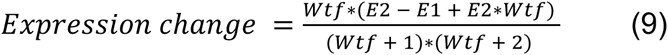

Solving gives us 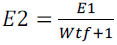 for which expression does not change. This is the upper limit of E2 for expression reduction, thus when E2 is below this limit we get expression reduction. At low [TF] (low Wtf) this limit is high, thus at low [TF] it’s easy to reduce expression. This limit goes to E2 = 0 when [TF] goes to infinity, thus at high [TF] there is no expression reduction. This is the behavior that we observe, suggesting that TF sharing is the mechanism behind the expression reduction as a result of activator site addition.

Taken together, we find that while both steric hindrance and TF sharing have regimes in which site addition decreases expression, only TF sharing shows a decrease when [TF] is low and less or no decrease when [TF] is high. In contrast, the steric hindrance model gives the opposite behavior and shows an increase in expression reduction as the [TF] goes up. Thus, a theoretic investigation of the effect of activator site addition on expression shows that TF sharing and not steric hindrance is, at least qualitatively, able to explain the expression reduction that we observe mostly at high [AA].

In order to investigate why adding a binding site can reduce expression we performed the following analysis. We measure the expression driven by the promoter with no Gcn4 binding sites (O), the two promoters each with a single Gcn4 binding site ( promoter A, promoter B), and the promoter with both binding sites ( promoter AB ). We observe that adding each binding sites separately results in an increase in expression compared to the no binding site promoter (A>O and B>O), suggesting that these binding sites, at least in isolation, do not recruit transcriptional repressors. Importantly, these two binding sites in isolation drive different expression levels, with A driving higher expression than B. However, we observe that the two Gcn4 binding sites together often results in lower expression than the highest of the single promoters (AB < max(A,B), and A>B)). The 'sharing' model suggests that the B (lower expression) binding site steals TF from A (the higher expression binding site), and therefore expression with AB is less than that from A (but still greater than expression from B). These results cannot be explained by additive repression.

### Supplementary data

Supplementary data are online at

https://www.upf.edu/scb/SupplementaryData/vanDijk_Sharon_TFSharing_2015.zip

**expression_all_constructs.tab**

Measured expression values of all constructs measured in the 6 conditions

**annotation_all_constructs.tab**

Sequence annotations for all promoters in the library

**\*_constructs_expression.tab**

Predicted expression values for model *, where * is: null (basic model), PSE (position specific expression), SH (steric hindrance), S (TF sharing), or a combination of these.

**\*_params.tab**

Model parameters for model *, where * is: null (basic model), PSE (position specific expression), SH (steric hindrance), S (TF sharing), or a combination of these.

